# Acute infectious mononucleosis generates persistent, functional EBNA-1 antibodies with high cross-reactivity to alpha crystalline beta

**DOI:** 10.1101/2024.12.18.629009

**Authors:** Krishna Kumar Ganta, Margaret McManus, Ross Blanc, Qixin Wang, Wonyeong Jung, Robin Brody, Mary Carrington, Robert Paris, Sumana Chandramouli, Ryan McNamara, Katherine Luzuriaga

## Abstract

Epstein-Barr Virus (EBV) infects over 95% of the world’s population and is the most common cause of infectious mononucleosis (IM). Epidemiologic studies have linked EBV with certain cancers or autoimmune conditions, including multiple sclerosis (MS). Recent studies suggest that molecular mimicry between EBV proteins, particularly EBV nuclear antigen 1 (EBNA-1), and self-proteins is a plausible mechanism through which EBV infection may contribute to the development of autoimmune disorders. We used a systems immunology approach to investigate the magnitude, specificity, and functional properties of EBNA-1 specific antibodies in a cohort of 97 young adults with IM from presentation through 1-year post-primary infection compared to a control cohort of EBV-seropositive individuals. Levels of EBNA-1 specific IgG1 and IgG3 binding antibodies increased over the course of infection. EBNA-1 antibodies capable of mediating antibody-dependent cellular phagocytosis (ADCP) and antibody-dependent complement deposition (ADCD) were detected at or after 6 months. Binding and ADCP- and ADCD-leveraged antibodies primarily targeted a region of EBNA-1 known to elicit cross-reactive antibodies to several self-peptides in individuals with MS. Significantly higher binding and ADCD-active antibodies targeting EBNA-1 were observed in individuals with at least one HLA-DRB1*15:01 allele, a known genetic risk factor for MS; Importantly, high levels of antibodies capable of binding alpha crystalline beta (CRYAB) and mediating complement deposition were detected at 6 months and 1-year following IM; CRYAB antibodies were resistant to denaturing forces, indicating an affinity matured response. Blocking experiments confirmed that CRYAB antibodies were cross-reactive with EBNA-1. Altogether, these results demonstrate that high levels of functional antibodies targeting EBNA-1 are generated in early EBV infection, some of which are cross-reactive with CRYAB. Further investigation is warranted to determine how these antibody responses may contribute to the subsequent development of MS.

## Introduction

Epstein-Barr Virus (EBV) establishes persistent infection in > 95% of the world’s population by the fourth decade of life^1^. Young children experience only mild or non-specific symptoms with primary EBV infection; however, primary EBV infection in older children, adolescents, and adults commonly causes infectious mononucleosis (IM)^1^. EBV has also been implicated in the development of various cancers and autoimmune disorders, most notably multiple sclerosis (MS) ^2,3^; for example, a history of symptomatic IM increases an individual’s risk of MS 2-3 fold ^4^. Evidence supporting a direct causal relationship between EBV and MS was provided by a recent study that compared antibody responses to the entire human virome in a large cohort of young military recruits who developed MS to those who did not develop MS ^3^. In this study, EBV infection, but not infection with any other virus, conferred an increased risk of developing MS; EBV infection acquired prior to 18 years of age conferred a 26-fold risk while infection later in life conferred a 32-fold risk of MS.

EBNA-1 is a DNA-binding protein necessary for replication and maintenance of the EBV genome in latently infected cells; it is the only EBV protein expressed in all EBV-associated cancers^5,6^. Multiple studies have linked high levels of EBNA-1 antibodies or T cell responses to a higher risk of development of autoimmune disorders ^3,7^. In particular, high levels of antibodies that target a region of EBNA-1 (aa 365-459) with known homology to several human proteins have been linked to an elevated risk of developing MS^3^; risk is especially high (15-fold) in individuals who have at least one HLA-DRB1*15:01allele and who are HLA-A*02:01 negative ^8^. A history of IM combined with high EBNA-1 antibody levels in individuals with HLA-DRB1*15:01 and who lack an HLA A*0201 allele further increases the risk (20-fold) of developing MS ^7^.

Several recent studies have demonstrated that sequence similarities between EBNA-1 peptides and self-peptides (“molecular mimicry”) result in the generation of autoreactive antibodies or T cell responses, providing a potential mechanism through which EBV may trigger autoimmune disorders. For example, several studies have reported the detection of EBNA-1 binding antibodies that cross-react with epitopes in alpha crystalline beta (CRYAB) ^9^, anoctamin 2 (ANO2) ^10^, or Glial Cell Adhesion Molecule (GlialCAM)^11^ in individuals with MS.

The present work evaluated EBNA-1-specific antibody magnitude, specificity, and function at presentation with and following acute IM in order to inform future work on how they might contribute to the pathogenesis of EBV-related autoimmune disorders. Through evaluation of antibody subclass, Fc receptor binding, and functions including phagocytosis, complement deposition, and cross-reactivity with self-peptides, we provide a deeper understanding of the evolution of EBNA-1 specific antibodies in primary EBV infection, as well as their potential role in the pathogenesis of subsequent autoimmune disorders.

## Results

### High levels of EBNA-1 IgG1 and IgG3 antibodies develop post IM and target an EBNA-1 region implicated in autoimmunity

Plasma samples were obtained from 97 young adults (median age 19 yrs) at IM presentation, and again at 6 weeks, 6 months, and 1-year post-IM presentation; 50 EBV seropositive young adults (median age 18.8 years); and 10 EBV seronegative young adults (Median age 18.5 yrs; Table 1). IM diagnosis was based on clinical symptoms and confirmatory serology, as previously described ^12,13^. Age, gender, race, and HLA types of the study cohort are provided in Table 1. When surveyed at the time of specimen collection, 20 EBV seropositive participants reported a history of IM; 30 EBV seropositive participants did not recall/did not report a history of IM.

We used a systems immunology approach, combining HLA typing with systems serology to comprehensively assess antibody responses. To achieve this, we used several peptides spanning the EBNA-1 protein, enabling a comprehensive evaluation of immune responses to different regions of the antigen (Figure S1). From the overall immune response heatmap (Figure S2), we observed that IgG1 and IgG3 responses were more pronounced compared to IgG2 and IgG4 across the EBV infection groups, justifying our focus on these isotypes to better understand their potential role in EBNA-1-specific immunity and autoimmune disease risk. This integrated framework enabled us to examine the interplay between genetic factors and antibody functionality, providing insights into how EBV infection and HLA alleles shape immune dynamics and contribute to autoimmune disease risk. EBNA-1 specific IgG1 and IgG3 antibodies were infrequently detected at presentation with IM; levels of these antibodies were significantly higher at 6 months and 1 year compared to levels at IM presentation (Figures 1A and 1C). At 1 year, EBNA-1 specific IgG1 levels did not differ significantly from those measured in EBV seropositive individuals. However, IgG3 levels were significantly higher at 1-year following IM than levels in seropositive individuals. EBNA-1 IgG1 and IgG3 antibodies primarily targeted peptides spanning amino acids 365-459 (Figure 1B and 1D). Antibodies to this segment of EBNA-1 have previously been identified to be elevated in individuals with MS, some of which are cross-reactive to peptides from human proteins ^9-11^. Interestingly, binding IgG1 levels to three EBNA-1 peptides (aa 365-420, aa 377-459, 393-448) were highly correlated with each other (R > 0.9, Figure S3), suggesting that these peptides target a common immunodominant epitope. Altogether, these data demonstrate that high levels of antibodies to an EBNA-1 region implicated in autoimmunity are commonly detected within 6 months of presentation with IM and persist beyond primary infection.

**Figure 1.**
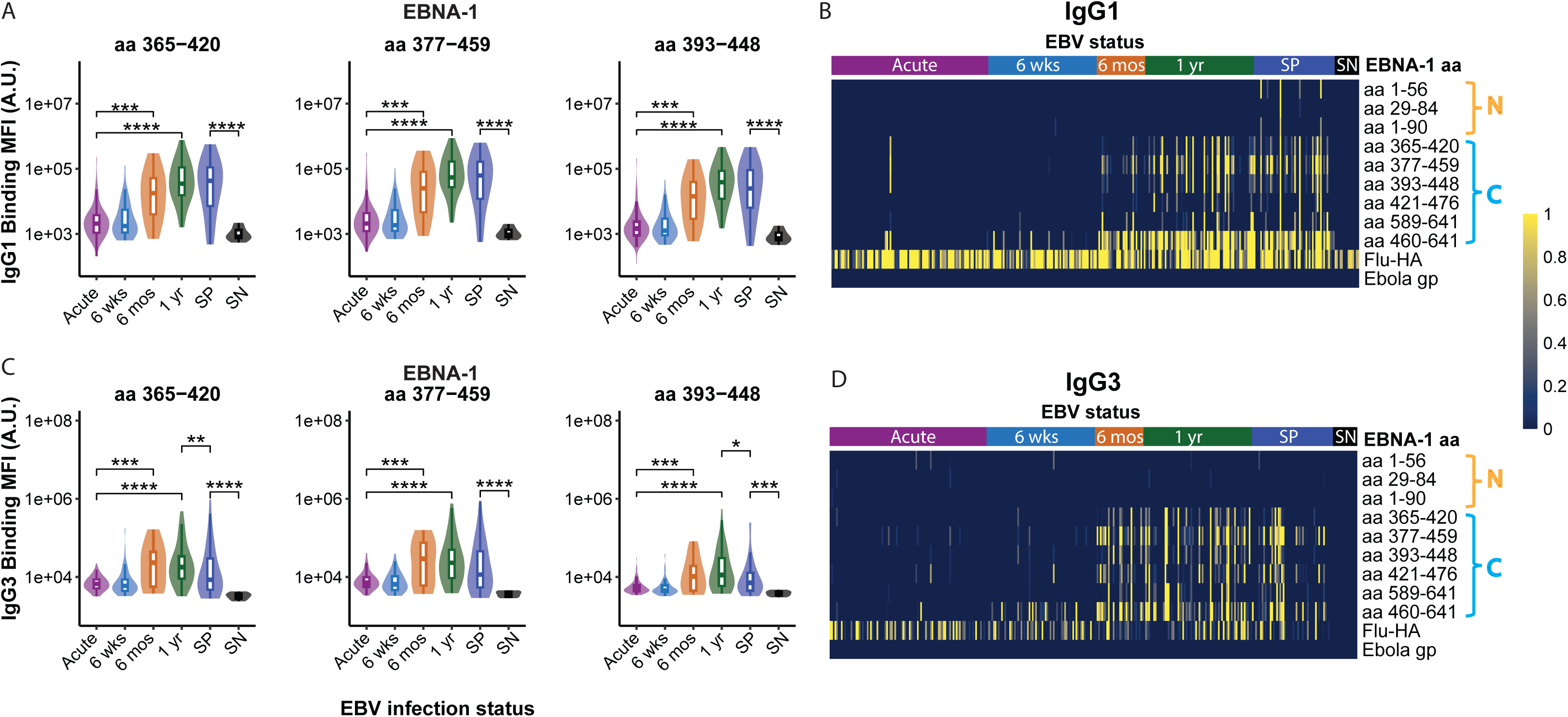
IgG1 and IgG3 Binding to EBNA-1 C-Terminal Peptides. (A) Violin and boxplot showing IgG1 responses towards EBNA-1 C-terminal domain peptides at: Acute presentation (n=97), 6 weeks (n=67), 6 months (n=30), and 1 year (n=67) post-IM diagnosis. EBV seropositive (SP, n=50) and EBV seronegative controls (SN, n=10) are also included. Y-axis units are IgG1 binding levels quantified through median fluorescence intensity (MFI) as arbitrary units (A.U.). (B) Overall IgG1 binding heatmap to regions of EBNA-1. Shown on the right-hand side are the peptides used. Influenza haemagglutinin (HA) is used as a positive control and Ebolavirus glycoprotein (GP) is used as a negative control. On top are the time points analyzed for the IM cohort, as well as SP and SN individuals as controls. Binding is shown as a fraction of maximum row binding, with the heatmap legend shown on the far right. (C) Same as A, but for IgG3 responses to EBNA-1 C-terminal domain peptides. (D) Same as B, but for IgG3 binding heatmap to regions of EBNA-1. For all comparisons, statistical significance is indicated as *p<0.05, **p<0.01, ***p<0.001, ***p<0.0001 using a Wilcoxon Test followed by Bonferroni correction for multiple comparisons. Non-significant differences were left blank.

### Higher levels of EBNA-1 binding antibodies are detected in individuals with HLA-DRB1*15:01

Individuals with a history of IM, high titers of an EBNA-1 region implicated in autoimmunity, and certain HLA alleles have a particularly elevated risk of MS ^7,8,14,15^. We therefore explored the relationship between binding EBNA-1 antibody levels and these reported risk factors. HLA-DRB1*15:01 is the strongest genetic factor associated with elevated MS risk ^8^. Approximately 20 % of IM individuals and seropositive controls in our cohort had at least 1 HLA-DRB1*15:01 allele (Table S1). IM individuals with at least one HLA-DRB1*15:01 allele had significantly higher IgG1 binding antibody levels targeting EBNA-1 peptides between aa 365-448 at 1-year post-infection compared to those without an HLA-DRB1*15:01 allele (Figure 2A). Individuals with IM and at least one HLA-DRB1*15:01 allele also had significantly higher fold expansion of IgG1 antibodies at 1-year relative to acute levels than those without an HLA-DRB1*15:01 allele (Figure 2B). A mixed model analysis confirmed a potential interaction between the EBV infection stage and HLA-DRB1*15:01 status. Trend plots (Figure S4) for EBNA-1 IgG1 antibody responses targeting the EBNA-1 region linked to autoimmunity demonstrated significantly higher antibody levels in individuals with HLA-DRB1*15:01 compared to those without the allele; this difference is most pronounced at 1-year post-infection. Individuals with at least one HLA-DQB1*06:02 or HLA-DQA1*01:02 allele also showed significantly higher fold changes from acute infection through convalescence, though their absolute antibody levels were not significantly different (Figure S5). Of note, the HLA-DQB1*06:02 and HLA-DQA1*01:02 alleles are in linkage disequilibrium with HLA-DRB1*15:01, and this haplotype confers an elevated risk of MS in European populations ^16^. Altogether, these results link the detection of high titers of antibodies targeting a segment of EBNA-1 that shares sequence homology (Figure S1) with human proteins with an HLA haplotype (HLA-DRB1*15:01, DQB1*06:02, DQA1*01:02) previously shown to be associated with MS risk.

**Figure 2.**
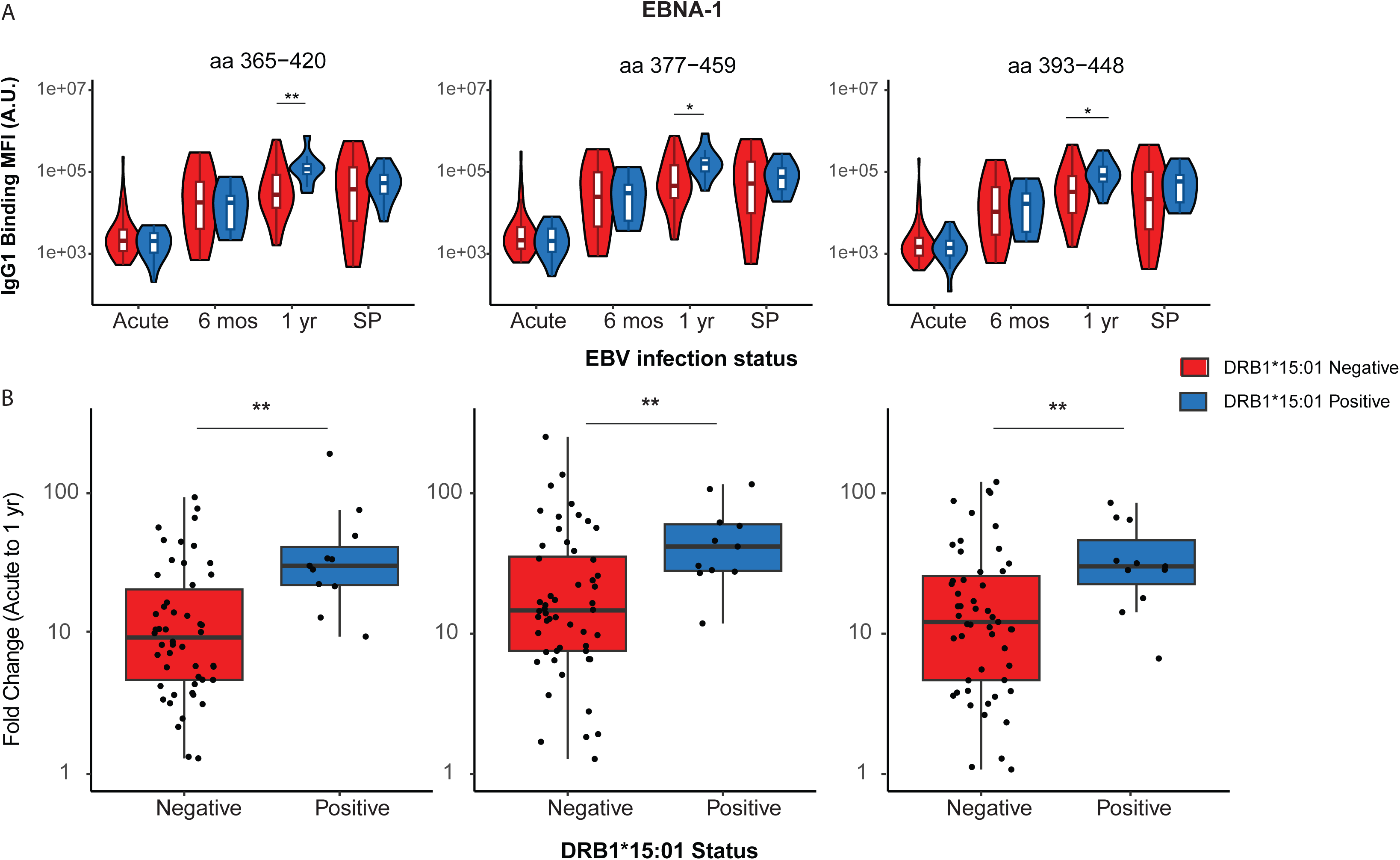
The presence of the HLA DRB1*15:01 allele enhances EBNA-1 IgG1 responses. (A) Violin and boxplot showing IgG1 responses towards EBNA-1 C-terminal domain peptides (aa 365-420, 377-459, 393-448) at acute, 6 months, and 1-year post-IM. Groups were distinguished based on being positive (blue) or negative (red) for at least 1 DRB1*15:01 allele. Shown for reference are SP individuals., and seropositive controls (SP) in individuals stratified by HLA DRB1*15:01 status. (B) Fold changes in IgG1 levels at 1-year post-IM relative to acute to EBNA-1 C-terminal domain peptides (aa 365-420, 377-459, 393-448) were plotted using box and whisker plots for individuals who were positive (blue) or negative (red) for at least one DRB1*15:01 allele. This comparison was made to control potential baseline/acute differences between the groups. The number of individuals in the IM cohort with at least one HLA-DRB1*15:01 allele: 19 of 97 participants (20%) at the acute visit; 8 of 30 (27%) participants at 6 months; and 11 of 67 (16%) participants at 1 year. In the SP individuals, there are 11 HLA-DRB1*15:01 positive and 38 HLA-DRB1*15:01 negative participants. For all comparisons, statistical significance is indicated as *p<0.05, **p<0.01, ***p<0.001, ***p<0.0001 using a Wilcoxon Test followed by Bonferroni correction for multiple comparisons. Non-significant differences were left blank.

### EBNA-1 C-terminal domain targeting antibodies are polyfunctional through FcγR-engagement

A growing body of evidence indicates that antibody engagement of effectors through their Fc domains is critical for mediating immune effector functions, including phagocytosis and complement activation, that are associated with infection outcomes^17^. We therefore measured the binding of EBNA-1 antibodies to Fc receptors. EBNA-1 antibodies capable of binding Fc receptors FcγRIIa and FcγRIIIa were detected at 6 months and 1 year and again primarily targeted peptides between aa 365-459 (Figure 3). Thus, the timing of the development of antibodies capable of engaging Fc receptors, as well as the antibody specificities, were similar to those observed for binding antibodies. This is at difference to some vaccine responses where binding antibodies persisted, but FcR-binding antibodies waned ^18^. This suggests a more durable, functionally active antibody response after IM.

**Figure 3.**
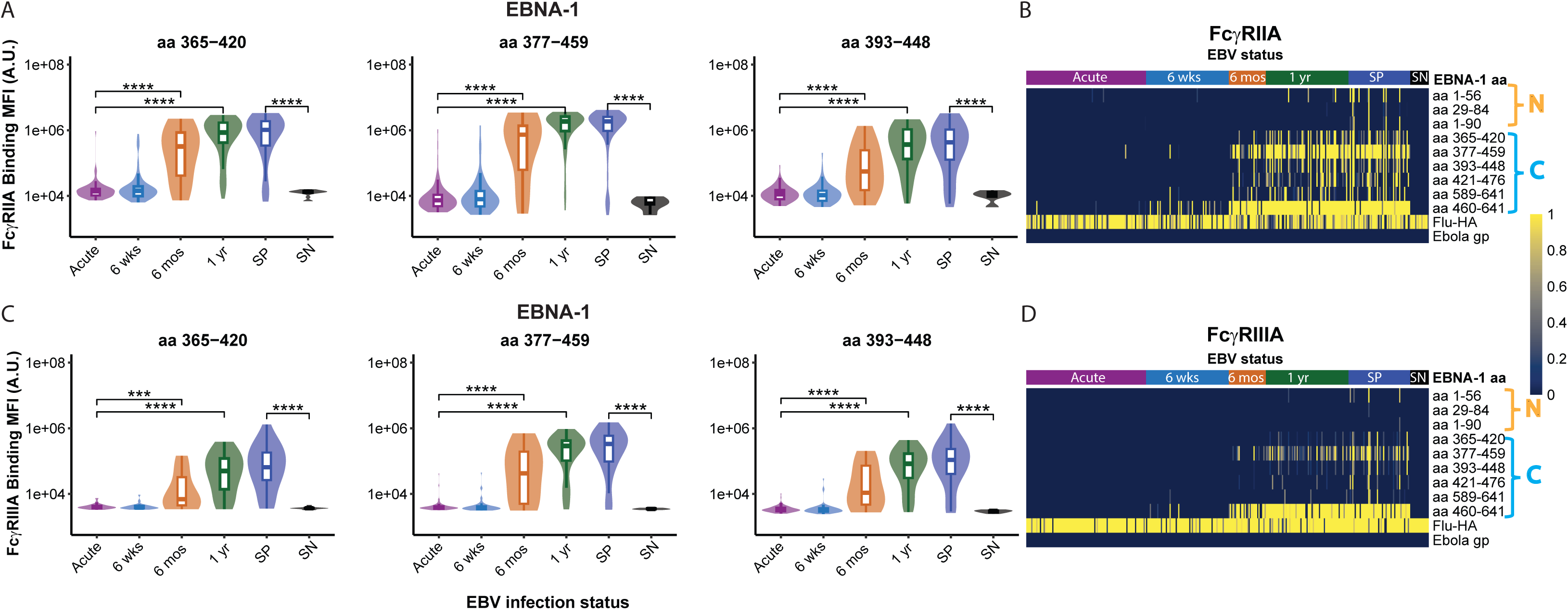
FcγRIIA and FcγRIIIA binding antibodies targeting EBNA-1 peptides persist up to 1-year post-IM. (A) Violin and boxplot showing FcγRIIA-binding antibody responses towards EBNA-1 C-terminal domain peptides at: Acute presentation (n=97), 6 weeks (n=67), 6 months (n=30), and 1 year (n=67) post-IM diagnosis. EBV seropositive (SP, n=50) and EBV seronegative (SN, n=10) are also included as controls. Y-axis units are FcγRIIA-binding antibody levels quantified through MFI as arbitrary units (A.U.). (B) Overall FcγRIIA-binding antibody heatmap to regions of EBNA-1. Shown on the right-hand side are the peptides used. Influenza HA (HA) is used as a positive control and Ebolavirus GP is used as a negative control. On top are the time points analyzed for the IM cohort, as well as SP and SN individuals as controls. Binding is shown as a fraction of maximum row binding, with the heatmap legend shown on the far right. (C) Same as A, but for FcγRIIIA-binding antibody responses towards EBNA-1 C-terminal domain peptides. (D) Same as C, but for FcγRIIIA-binding antibody heatmap to regions of EBNA-1. For all comparisons, statistical significance is indicated as *p<0.05, **p<0.01, ***p<0.001, ***p<0.0001 using a Wilcoxon Test followed by Bonferroni correction for multiple comparisons. Non-significant differences were left blank.

We evaluated the functional capacity of antibodies targeting EBNA-1 peptides using an assay that measured phagocytosis of immune complexes composed of plasma antibodies bound to EBNA-1 peptide (aa 365-420, 377-459, aa 393-448) coated beads. Plasma ADCP activity at the 6-month and 1-year visits differed significantly from ADCP activity in plasma samples from the acute visit. (Figure 4); plasma ADCP activity was also detected in plasma samples from EBV seropositive individuals. This is consistent with the persistently high FcγR-binding antibodies in this cohort.

**Figure 4.**
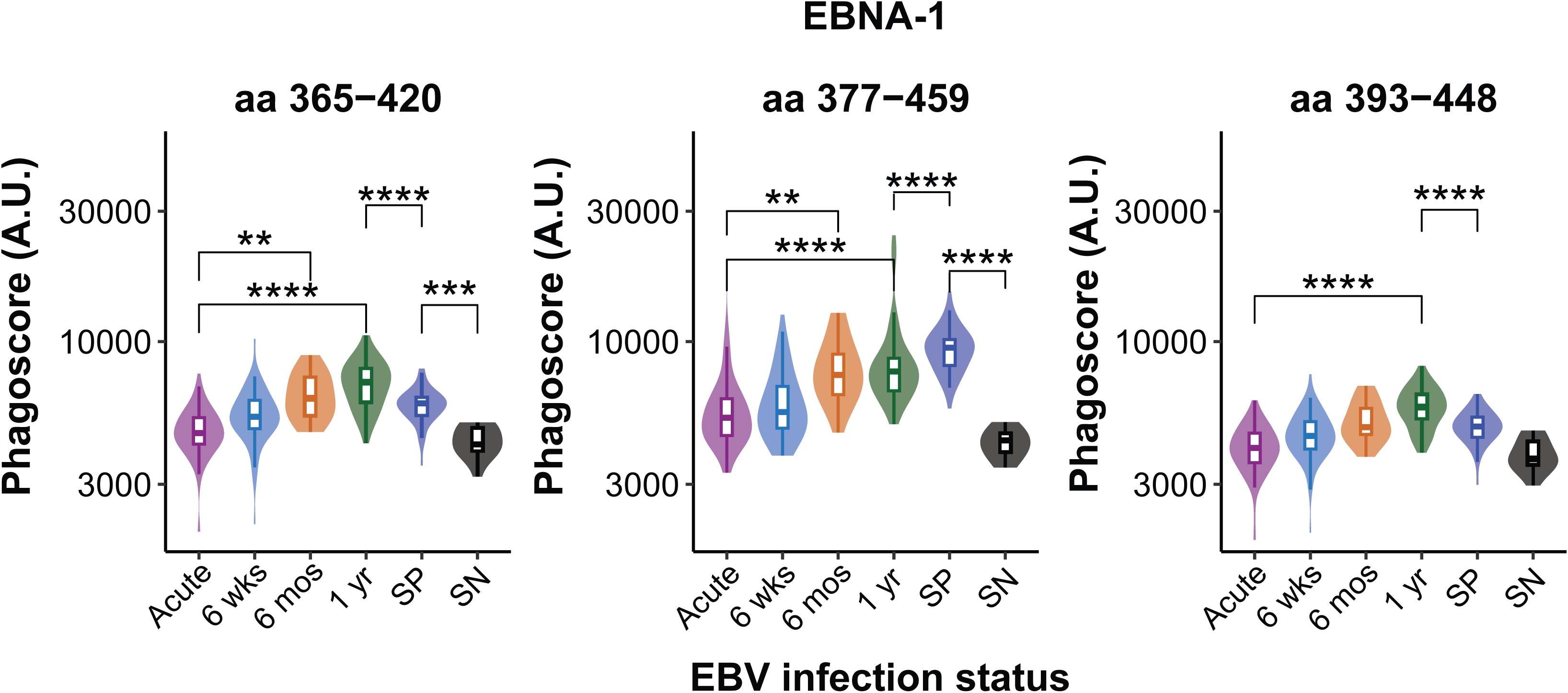
Antibody-dependent cellular phagocytosis is activated post-IM to the EBNA-1 C-terminal domain. Violin and boxplots showing antibody-dependent cellular phagocytosis by monocytes (ADCP) quantified through phagoscores (see methods). Phagoscores were quantified against EBNA-1 peptides (aa 377-459, 365-420, and 393-448) across different EBV infection stages: Acute, 6 weeks, 6 months, and 1-year post-IM. EBV seropositive (SP) and seronegative (SN) are included as controls. For all comparisons, statistical significance is indicated as *p<0.05, **p<0.01, ***p<0.001, ***p<0.0001 using a Wilcoxon Test followed by Bonferroni correction for multiple comparisons. Non-significant differences were left blank.

### EBNA-1 antibodies mediate complement activation and deposition

Having measured high levels of EBNA-1 binding antibodies and Fc receptor binding, we next evaluated the capacity of these antibodies to activate and fix complement. Antibodies targeting EBNA-1 peptides and capable of fixing complement were detected at significantly higher levels at 6 months and 1 year than at IM presentation (Figures 5A and 5B). A mixed-model analysis (Figure 5C) revealed that by 1-year post-infection, individuals with at least one DRB1*1501 allele had significantly higher ADCD activity compared to individuals without a DRB1*1501 allele in response to EBNA-1 C-terminal peptides 365-420 (p < 0.001) and 377-459 (p < 0.001).

**Figure 5.**
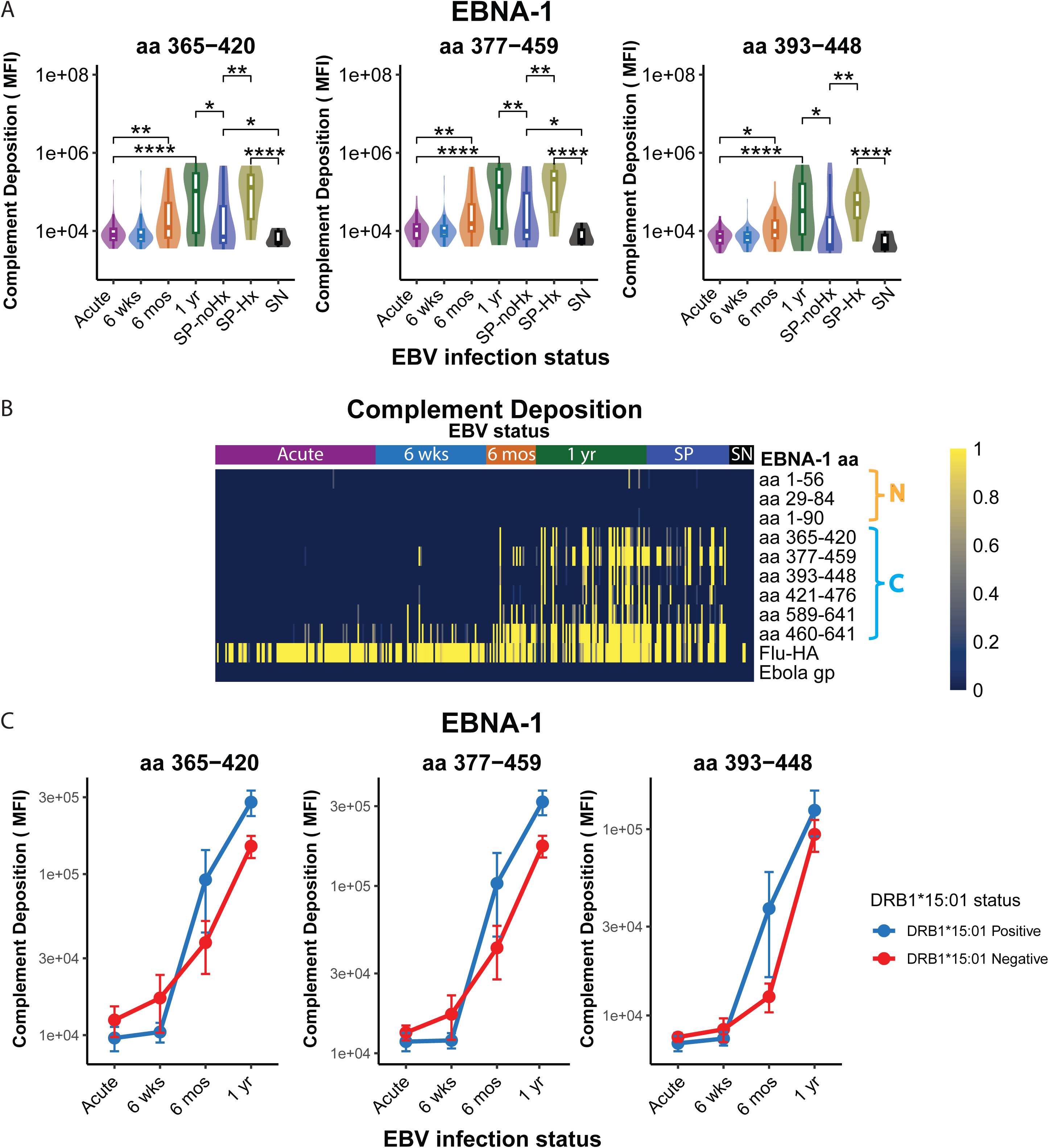
Antibody-dependent complement deposition (ADCD) and mixed model analysis of EBNA-1 ADCD activity to C-terminal peptides. (A) Violin and boxplots showing antibody-dependent complement deposition (ADCD) quantified through C3 deposition units for EBNA-1 peptides (aa 377-459, 365-420, 393-448) across) across different EBV infection stages: Acute, 6 weeks, 6 months, and 1-year post-IM. EBV seropositive (SP) and seronegative (SN) are included as controls. For all comparisons, statistical significance is indicated as *p<0.05, **p<0.01, ***p<0.001, ***p<0.0001 using a Wilcoxon Test followed by Bonferroni correction for multiple comparisons. Non-significant differences were left blank. (B) Overall ADCD to regions of EBNA-1. Shown on the right-hand side are the peptides used. Influenza HA is used as a positive control and Ebolavirus GP is used as a negative control. On top are the time points analyzed for the IM cohort, as well as SP and SN individuals as controls. The scale is shown as a fraction of the maximum row ADCD, with the heatmap legend shown on the far right. (C) Line graphs showing ADCD over time to EBNA-1 peptides (aa 377-459, 365-420, 393-448) for individuals who were positive (blue) or negative (red) for at least one DRB1*15:01 allele. Statistical analysis was performed using a linear mixed model with interaction terms for EBV infection stage and DRB1*15:01 status. Significant increases in ADCD were observed at 1 year compared to Acute, 6 weeks, and 6 months for all peptides (p < 0.001). Interaction effects at the 1-year time point revealed that DRB1*15:01 positive individuals showed a more pronounced ADCD response compared to negatives for the peptides aa 377-459 (p = 0.000737) and 365-420 (p = 0.000875). No significant interaction effects were observed at earlier time points.

Complement-fixing EBNA-1 antibodies were also detected in EBV seropositive individuals. On a questionnaire completed at the first study visit, 30 EBV seropositive individuals in our cohort did not recall a history of IM while 20 seropositive individuals reported a prior diagnosis of IM. Significantly higher levels of EBNA-1 antibodies capable of fixing complement were detected in individuals who related a history of IM compared to individuals without a history of IM (Figure 5A).

### High levels of binding and ADCD antibodies to CRYAB are detected following IM

We next evaluated whether individuals with IM develop antibody responses to self-peptides (GlialCAM, anoctamin 2, and CRYAB) that share sequence homology with EBNA-1 (Figure S1), as described in individuals with MS. Figure 6A shows that binding IgG antibodies targeting CRYAB were detected at 6 months and 1 year. The detection of binding antibodies to self-proteins in our IM cohort is compatible with at least 2 reports of transient detection of antibodies to self-peptides in IM ^19,20^; in our study, binding IgG antibodies to CRYAB were not transient but persisted through 1-year and were also detected in EBV seropositive individuals. Of note, we did not detect antibodies to CRYAB in EBV seronegative individuals, nor did we detect antibodies to anoctamin 2 or GlialCAM in any individuals at or after presentation with IM (Figure S6) We next asked if these cross-reactive antibodies were weakly binding and could be dislodged with the disrupting agent urea. This approach has been previously used to distinguish avid antibodies from non-specific antibodies ^21-23^. Interestingly, antibody binding was resistant to dissociation in the presence of urea (Figure 6B), implying that the CRYAB antibodies detected at 6 months and 1 year following IM, as well as those detected in EBV seropositive individuals, were of high avidity.

**Figure 6.**
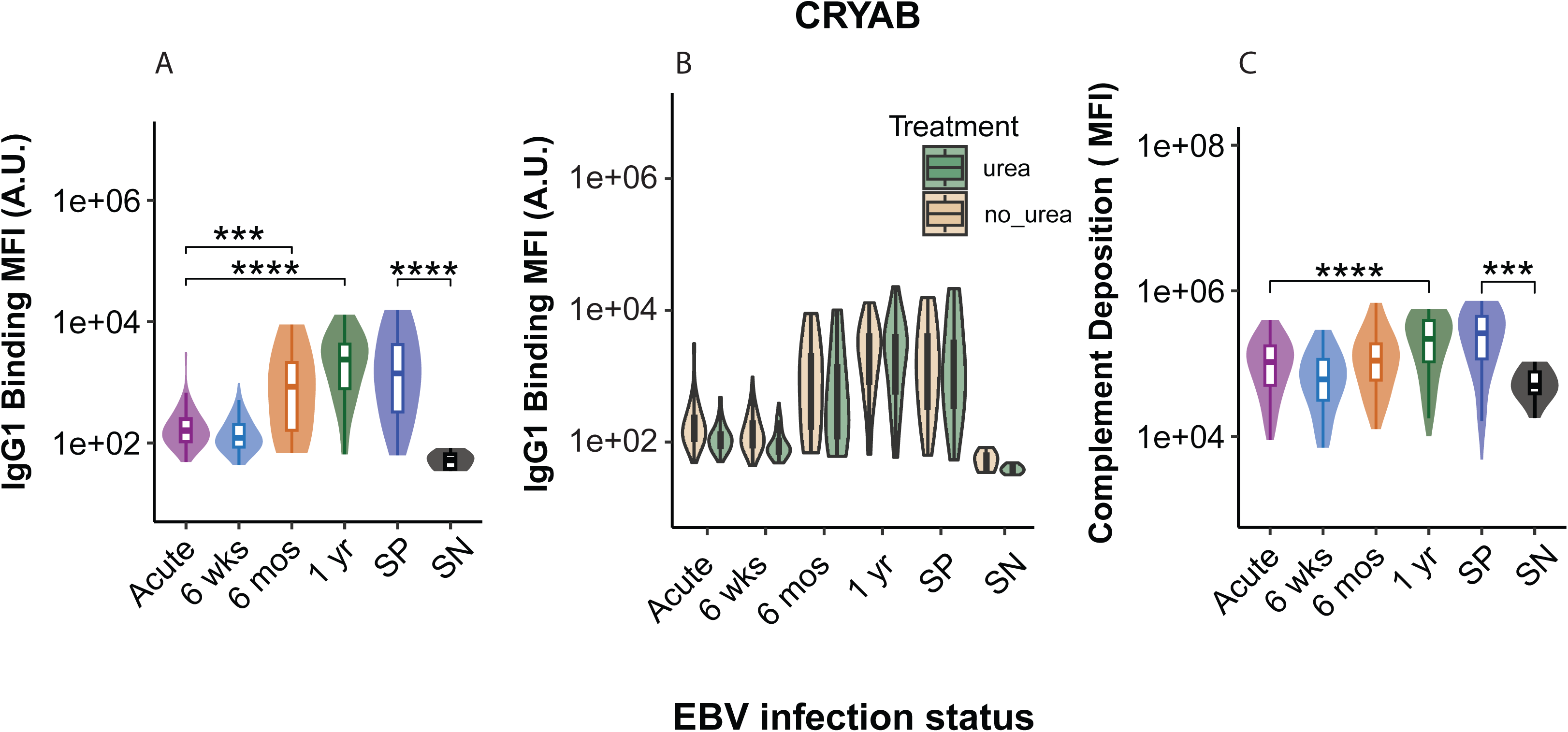
EBNA-1 antibodies are cross-reactive to CRYAB and retain avidity and ADCD to the self-peptide. (A) Violin and boxplots showing IgG1 binding to CRYAB across different EBV infection stages: Acute, 6 weeks, 6 months, and 1-year post-IM. EBV seropositive (SP) and seronegative (SN) are included as controls. (B) Violin and boxplots showing IgG1 binding to CRYAB in the presence (green) or absence (brown) of 3M urea across different EBV infection stages: Acute, 6 weeks, 6 months, and 1-year post-IM. EBV seropositive (SP) and seronegative (SN) are included as controls. (C) Violin and boxplots showing ADCD to CRYAB across different EBV infection stages: Acute, 6 weeks, 6 months, and 1-year post-IM. EBV seropositive (SP) and seronegative (SN) are included as controls. For all comparisons, statistical significance is indicated as *p<0.05, **p<0.01, ***p<0.001, ***p<0.0001 using a Wilcoxon Test followed by Bonferroni correction for multiple comparisons. Non-significant differences were left blank.

The complement system is increasingly recognized for its role in the pathogenesis of neuroinflammatory and neurodegenerative diseases, including MS^24^. We therefore went on to evaluate whether the CRYAB IgG antibodies detected in binding assays were capable of mediating ADCD. We observed a significant increase in ADCD activity to CRYAB from the acute phase of IM through 6 months and 1 year; we also detected CRYAB-specific ADCD in seropositive individuals (Figure 6) but not in EBV seronegative individuals.

### CRYAB antibodies are cross-reactive with EBNA-1 antibodies

To further test whether EBNA-1 antibodies could bind to CRYAB, we set up an antigen-specific antibody depletion assay using a peptide blocking approach ^9^. Serum samples were spiked with EBNA-1 peptides to deplete EBNA-1-specific antibodies, and their cross-reactivity with CRYAB was assessed. If EBNA-1 antibodies are cross-reactive, their depletion would result in a lack of detectable binding to CRYAB. To validate the specificity of this approach, we performed a control blockade by spiking the same samples with flu peptides to ensure that flu-specific antibodies do not interfere with anti-CRYAB antibody levels. Additionally, VCA IgG levels were also measured under both blockade conditions to confirm that the depletion is specific to EBNA-1 or flu antibodies without affecting other antibody populations.

Anti-CRYAB IgG levels were significantly (>90%) reduced when plasma samples from convalescent and seropositive individuals were depleted with EBNA-1 peptides (365-420,377-449, and 393-449), indicating a specific cross-reactivity between CRYAB antibodies and EBNA-1 (Figure 7). By contrast, treatment with Flu-HA peptide did not reduce CRYAB total IgG levels, confirming the specificity of this cross-reactivity. Anti-Flu-HA total IgG reactivity was reduced by Flu-HA peptide but not by EBNA-1 peptides; anti-VCA IgG levels remained unchanged under both blocking conditions, demonstrating the specificity of the observed CRYAB and EBNA-1 cross-reactivity. Altogether, these results confirm the generation of CRYAB and EBNA-1 cross-reactive antibodies shortly following primary EBV infection.

**Figure 7.**
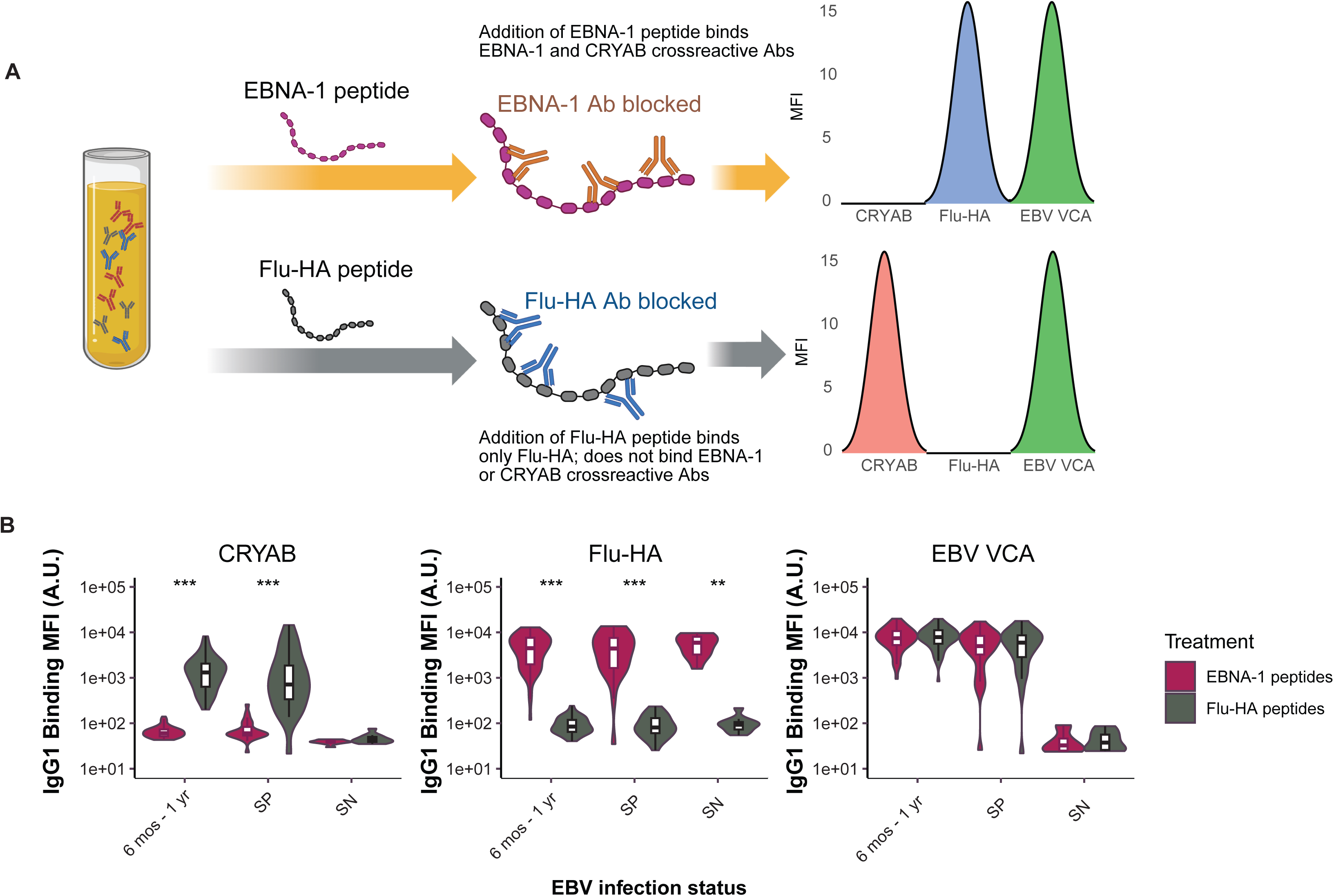
CRYAB autoantibodies from post-IM individuals are specific to EBNA-1 responses. (A) Schematic of the blocking assay: EBNA-1 peptides were used to deplete immunoglobulins targeting the viral protein. If these antibodies are cross-reactive with CRYAB, then the EBNA-1 immunodepletion should result in a loss of CRYAB binding. As a control, the same serum was blocked with Flu-HA peptides to control non-specific immunodepletion. Lastly, responses to VCA were quantified for both EBNA-1 and Flu-HA immunodepletions as this is not cross-reactive with either or shouldn’t result in any binding loss. (B) Violin and boxplots depicting IgG1 antibody reactivity to CRYAB, Flu-HA, and EBV VCA across different treatments (EBNA-1 peptides or Flu-HA peptides). Samples included: pooled 6 mos -1 yr (n = 55), EBV seropositive (SP, n = 25), and EBV seronegative (SN, n=10). For all comparisons, statistical significance is indicated as *p<0.05, **p<0.01, ***p<0.001, ***p<0.0001 using a Wilcoxon Test followed by Bonferroni correction for multiple comparisons. Non-significant differences were left blank.

## Discussion

Multiple epidemiologic studies over the years have linked EBV infection to the risk of developing MS ^2,25,26^. A history of symptomatic IM confers additional risk (2-3-fold) ^4,25^. Molecular mimicry between EBNA-1 and self-peptides has been proposed as a plausible mechanism ^9-11,20^. We thus undertook this study which aimed to characterize the development of EBNA-1 specific antibody responses post-IM, and to determine whether cross-reactive antibodies are generated following primary EBV infection. High levels of IgG1, IgG3, and FcγR-binding antibodies were detected against the EBNA-1 C-terminal domain at 6 months and 1-year following IM (Figures 1-3). EBNA-1 specific antibodies demonstrated strong functional activity, including antibody-mediated phagocytosis and complement deposition (Figures 4 and 5). Levels of binding and complement-fixing EBNA-1 antibodies were higher in individuals with the MS-associated HLA-DRB1*15:01 allele than in individuals without HLA-DRB1*1501 (Figures 2 and 5). These EBNA-1 C-terminal domain antibodies showed high cross-reactivity and avidity to CRYAB, providing a direct link between EBV infection and the development of autoreactive antibodies to CRYAB (Figures 6 and 7). Other self-targeting antibodies such as those to ANO2 and GlialCAM were not detected through our method, indicating that anti-CRYAB development post-IM was specific, at least in the timeframe analyzed. Previous work has shown that autoantibodies to CRYAB were associated with MS^9^. Our work extends these studies by demonstrating that CRYAB autoantibodies generated post-IM are detectable in as little as 6 months after infection, are highly leveraged for effector function, and are linked to EBNA-1 C-terminal domain targeting antibodies.

EBNA-1 plays a key role in EBV persistence within latently infected B cells ^5^ and elicits antibody responses in most individuals with EBV infection. Prior studies have identified a region of EBNA-1 associated with autoimmune diseases ^15^. For example, several studies have documented elevated levels of binding antibodies specifically targeting EBNA-1 aa 365-448 or EBNA-1 377-459 in individuals with MS ^3,27^. A recent case-control study of military recruits described an elevated risk of developing MS in individuals with high EBNA-1 antibodies to C-terminal peptides (particularly aa 365-420). In these individuals, elevated levels of serum neurofilament light chain, a marker for neuronal damage, were detected a median of 6 years prior to MS onset ^3^. In our study, we demonstrate a significant increase in the levels of antibodies targeting these EBNA-1 peptides within 6 months of IM.

Significantly higher levels of IgG1 binding antibodies against C-terminal EBNA-1 peptides were detected in individuals carrying at least one HLA-DRB1*15:01 allele, which has been identified as the strongest identified genetic risk factor for MS ^28^. Our data are compatible with prior studies demonstrating an association between elevated binding antibody responses to EBNA-1 and HLA-DRB1*15:01^7,15^. We further add to these findings by demonstrating that these responses are not limited to individuals already diagnosed with MS but are also present in individuals shortly after primary EBV infection (Figure 2). Interestingly, we also observed that individuals carrying alleles (DQB1*06:02, DQA1*01:02) in linkage disequilibrium with HLA DRB1*1501^29^, exhibited significantly higher fold changes in EBNA-1 antibody levels over time (Figure S5). This suggests that this haplotype influences antibody production and function in ways that warrant further investigation.

Many individuals with elevated EBNA-1 IgG binding antibody titers also had antibodies capable of mediating phagocytosis and complement deposition (Figures 4 and 5). However, we also identified individuals with lower binding antibody levels and high functional activity, implying that antibody efficiency can compensate for lower titers^30^, likely driven by the antibody’s Fc region and its ability to engage Fc receptors ^17,31^. This finding underscores the importance of measuring both binding and functional activity to fully understand the mechanisms through which antibodies may be active in disease pathogenesis. Our mixed-model analysis revealed that individuals with the HLA-DRB1*15:01allele showed significantly higher levels of complement deposition activity at 1-year post-infection, particularly in response to EBNA-1 peptides within amino acids 365-420 and 377-459 (Figure 5C). While complement activation can be important for pathogen clearance, its dysregulation is also implicated in immunopathology. ^32^. Studies of post-mortem brain tissue from individuals with MS have consistently shown complement deposition in white matter plaques and gray matter lesions, suggesting that complement activation plays a role in disease progression^33^. In our study, the strong complement activation mediated by EBNA-1 specific antibodies, particularly in DRB1*1501 positive individuals, suggests that these antibodies could contribute to complement-driven damage observed in MS. Moreover, our results strongly support that the complement deposition was the result of affinity-matured antibody responses and not acute-phase responses dominated by IgM.

A key and novel aspect of our study was the detection of antibodies cross-reactive between EBNA-1 and CRYAB, a protein expressed in oligodendrocytes ^34^ cells within the central nervous system responsible for the production of myelin. Prior studies have demonstrated the presence of EBNA-1 antibodies cross-reactive with CRYAB in individuals with MS ^9^. In the present study, we show that the generation of EBNA-1 and CRYAB cross-reactive antibodies occurs in many individuals shortly after primary EBV infection; these cross-reactive antibodies also activate complement. While IgM can mediate complement deposition, the urea dissociation assays imply that affinity-matured IgG responses, rather than IgM, are responsible for the complement-fixing activity. The fact that these cross-reactive antibodies also activate complement adds to the concern that they could contribute to inflammatory processes and demyelination in the brain, potentially setting the stage for MS. However, given the rarity of MS relative to how commonly we detected EBNA-1, CRYAB, and cross-reactive antibodies following IM and in EBV seropositive individuals, additional factors are also likely involved in MS onset and disease progression.

Our study has several limitations that must be acknowledged. First and foremost, the follow-up period in this study was limited to 1-year post-infection. As a result, we are unable to determine whether the elevated antibody responses and functional activities observed, particularly in individuals with the DRB1*1501 allele, persist beyond this timeframe or whether they are associated long-term with the development of MS. Retrospective analysis of longitudinal samples from individuals with MS, as well as long-term follow-up of individuals experiencing IM may provide insights on whether these early immune responses are predictive of future disease risk. The history of IM in seropositive individuals was self-reported and medical records were not available to confirm participant reports.

Additionally, while we focused on the antibody responses and their functional capacities, other aspects of the immune response, such as T cell responses, were not evaluated. Future research should integrate these immune factors to provide a more comprehensive, systems-level understanding of the immune response to EBV infection in IM and its role in autoimmunity.

In conclusion, our study documents the early generation of high levels of antibodies targeting the C-terminal of EBNA-1 following IM. Levels of EBNA-1 binding and complement-fixing antibodies were highest in individuals with the HLA-DRB1*15:01 allele. The sustained antibody responses, their ability to activate complement, and their cross-reactivity with a central nervous system protein, CRYAB, suggest that EBNA-1 specific antibodies generated in early infection may play a role in the initiation of MS. Additional studies defining whether EBNA-1 antibody specificity and function can aid in early identification of individuals at risk of developing MS, as well as their potential role in the development of disease, could inform the development of strategies for early diagnosis and therapeutic intervention.

## ACKNOWLEDGEMENTS

We would like to extend our gratitude to the individuals who participated in these studies. We thank Yuko Yuki for HLA typing.

Funding for this project was provided by Moderna Therapeutics to RPM and KL; clinical research resources were provided by the UMass Center for Clinical and Translational Science (NIH UL1TR001453). This project has been funded in whole or in part with federal funds from the Frederick National Laboratory for Cancer Research, under Contract No. 75N91019D00024. The content of this publication does not necessarily reflect the views or policies of the Department of Health and Human Services, nor does mention of trade names, commercial products, or organizations imply endorsement by the U.S. Government. This Research was supported in part by the Intramural Research Program of the NIH, Frederick National Lab, Center for Cancer Research.

---

**Table S1.** Participant Characteristics.

|  | AIM | SeroPositive | SeroNegative |
| --- | --- | --- | --- |
| Number | 97 | 50 | 10 |
| Female sex (%) | 57 (59%) | 43 (86%) | 9 (90%) |
| Age in Years<br>(median,IQR) | 19.4 (18.0 – 22.6) | 18.8 (18.1 – 21.0) | 19.1 (18.4 – 22.6) |
| Race and Ethnicity (%) |  |  |  |
| White, Non-Hispanic | 89 (92%) | 35 (70%) | 9 (90%) |
| Black, Non-Hispanic | 1 (1%) | 2 (4%) | 0 (0%) |
| Asian, Non-Hispanic | 3 (3%) | 8 (16%) | 0 (0%) |
| Hispanic or Latino | 1 (1%) | 4 (8%) | 0 (0%) |
| Other, Non-Hispanic | 3 (3%) | 1 (2%) | 1 (10%) |
| HLA |  |  |  |
| A*0201 positive | 44 (45%) | 21 (42%) | 1 (14%)* |
| DRB1*1501 positive | 19 (20%) | 11 (22%) | 1 (14%)* |
| DQA1*0102 | 19 (20%) | 11 (22%) | 1 (14%)* |
| DQB1*0602 | 19 (20%) | 11 (22%) | 1 (14%)* |

\* HLA typing available for 7 of the 10 Seronegative participants

## STAR METHODS

### Key resource Table

| REAGENT | SOURCE | IDENTIFIER |
| --- | --- | --- |
| **Antibodies** |  |  |
| Anti-guinea pig complement C3 goat IgG fraction, FITC | MP Biomedicals | Cat # 0855385 |
| Anti-human IgG1 PE [HP6001] | Southern Biotech | Cat #9054-09 |
| Anti-human IgG3 PE [HP6050] | Southern Biotech | Cat #9210-09 |
| **Biological samples** |  |  |
| LowTox Guinea Pig Complement | CedarLane Labs | Cat # CL4051 |
| **Chemicals, peptides, and recombinant proteins** |  |  |
| Human soluble FcγR2A | Duke University | Custom Order |
| Human soluble FcγR3A | Duke University | Custom Order |
| LC-LC-Sulfo-NHS Biotin | ThermoFisher | Cat # A35358 |
| Streptavidin-R-PE | Prozyme | Cat # PJ31S |
| **EBNA-1 peptides** |  |  |
| Recombinant Epstein-Barr virus 1-90 | SinoBiological | Custom Synthesis |
| Recombinant Epstein-Barr virus 377-459 | SinoBiological | Custom Synthesis |
| Recombinant Epstein-Barr virus 460-641 | SinoBiological | Custom Synthesis |
| Recombinant Epstein-Barr virus 29-84 | SinoBiological | Custom Synthesis |
| Recombinant Epstein-Barr virus 365-420 B95-8 | SinoBiological | Custom Synthesis |
| Recombinant Epstein-Barr virus 393-448 B95-8 | SinoBiological | Custom Synthesis |
| Recombinant Epstein-Barr virus 393-448 AG876 | SinoBiological | Custom Synthesis |
| Recombinant Epstein-Barr virus EBNA-1 421-476 B95-8 | SinoBiological | Custom Synthesis |
| Recombinant Epstein-Barr virus EBNA-1 589-641 B95-8 | SinoBiological | Custom Synthesis |
| Recombinant Epstein-Barr virus EBNA-1 589-641 AG876 | SinoBiological | Custom Synthesis |
| Recombinant Epstein-Barr virus EBNA-1 1-56 GD1 | SinoBiological | Custom Synthesis |
| Recombinant Epstein-Barr virus EBNA-1 1-56 B95-8 | SinoBiological | Custom Synthesis |
| Recombinant Epstein-Barr virus EBNA-1 421-476 AG876 | SinoBiological | Custom Synthesis |
| Recombinant Epstein-Barr virus EBNA-1 449-504n B95-8 | SinoBiological | Custom Synthesis |

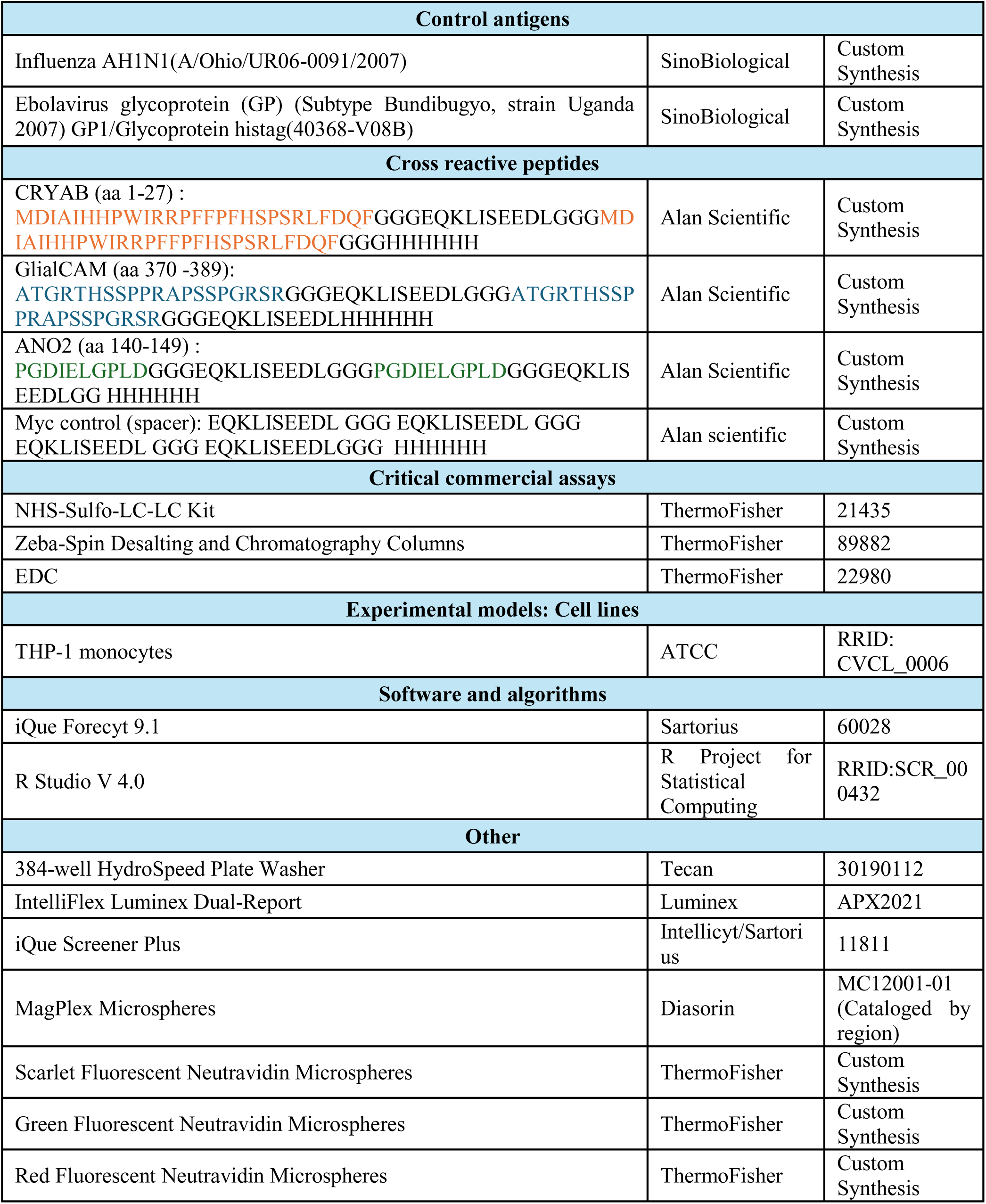

| Control antigens |  |  |
| --- | --- | --- |
| Influenza AH1N1(A/Ohio/UR06-0091/2007) | SinoBiological | Custom Synthesis |
| Ebolavirus glycoprotein (GP) (Subtype Bundibugyo, strain Uganda 2007) GP1/Glycoprotein histag(40368-V08B) | SinoBiological | Custom Synthesis |
| Cross reactive peptides |  |  |
| CRYAB (aa 1-27) :<br>MDIAIHHPWIRRPFFPFHSPSRLFDQFGGGEQKLISEEDLGGGMD<br>IAIHHPWIRRPFFPFHSPSRLFDQFGGGHHHHHH | Alan Scientific | Custom Synthesis |
| GlialCAM (aa 370 -389):<br>ATGRTHSSPPRAPSSPGRSRGGGEQKLISEEDLGGGATGRTHSSP<br>PRAPSSPGRSRGGGEQKLISEEDLHHHHHH | Alan Scientific | Custom Synthesis |
| ANO2 (aa 140-149) :<br>PGDIELGPLDGGGEQKLISEEDLGGGPGDIELGPLDGGGEQKLIS<br>EEDLGG HHHHHH | Alan Scientific | Custom Synthesis |
| Myc control (spacer): EQKLISEEDL GGG EQKLISEEDL GGG<br>EQKLISEEDL GGG EQKLISEEDLGGG HHHHHH | Alan scientific | Custom Synthesis |
| Critical commercial assays |  |  |
| NHS-Sulfo-LC-LC Kit | ThermoFisher | 21435 |
| Zeba-Spin Desalting and Chromatography Columns | ThermoFisher | 89882 |
| EDC | ThermoFisher | 22980 |
| Experimental models: Cell lines |  |  |
| THP-1 monocytes | ATCC | RRID:<br>CVCL_0006 |
| Software and algorithms |  |  |
| iQue Forecyt 9.1 | Sartorius | 60028 |
| R Studio V 4.0 | R Project for<br>Statistical<br>Computing | RRID:SCR_00<br>0432 |
| Other |  |  |
| 384-well HydroSpeed Plate Washer | Tecan | 30190112 |
| IntelliFlex Luminex Dual-Report | Luminex | APX2021 |
| iQue Screener Plus | Intellicyt/Sartori<br>us | 11811 |
| MagPlex Microspheres | Diasorin | MC12001-01<br>(Cataloged by<br>region) |
| Scarlet Fluorescent Neutravidin Microspheres | ThermoFisher | Custom<br>Synthesis |
| Green Fluorescent Neutravidin Microspheres | ThermoFisher | Custom<br>Synthesis |
| Red Fluorescent Neutravidin Microspheres | ThermoFisher | Custom<br>Synthesis |

### Lead contact

(Katherine Luzuriaga)

### Material availability

This study did not generate new unique reagents.

### Data and code availability

The dataset (Luminex and flow cytometry MFI values) generated for the study have been deposited on GitHub (Accession code: HSPHSystemsSerology/ProjID_KG20241203) They are publicly available from the date of publication.

This paper does not report the original code.

Any additional information required to reanalyze the data reported in this paper is available from the lead contact upon request.

### Material and Methods

#### IM cohort

Plasma samples were obtained from 97 young adults (median age 19 yrs) at IM presentation, and again at 6 weeks (N=67), 6 months (N=30), and 1-year post-IM presentation (N=67); 50 EBV seropositive young adults (median age 18.8 years); and 10 EBV seronegative young adults (Median age 18.5 yrs.) IM diagnosis was based on clinical symptoms and confirmatory serology, as previously described ^12,13^. Age, gender, race, and HLA types of the study cohort are provided in Table S1. When surveyed at the time of specimen collection, 20 EBV seropositive participants reported a history of IM; 30 EBV seropositive participants did not recall/did not report a history of IM.

### Ethical Considerations

This project was approved by the Institutional Review Boards (IRBs) at UMass Chan Medical School and Massachusetts General Hospital. Written informed consent was obtained from all participants.

### HLA typing

HLA class I and II genotyping was performed using a targeted next-generation sequencing method as described previously (1) with some modifications. Locus-specific primers were used to amplify 25 polymorphic exons of HLA-A and -B (exons 1–4), HLA-C (exons 1–5), HLA-E (exon 3), DPA1 (exon 2), DPB1 (exons 2–4), DQA1 (exons 1–3), DQB1 (exons 2 and 3), DRB1 (exons 2 and 3), and DRB3, 4, 5 (exon 2) genes with Fluidigm Access Array System and Juno LP 48.48 IFCs (Fluidigm). The 25 Fluidigm PCR amplicons were pooled and subjected to sequencing on an Illumina MiSeq sequencer (Illumina). HLA alleles and genotypes were called using the Omixon HLA Explore (version 2.0.0) software (Omixon) ^35^.

### Antigens

EBNA-1 peptides spanning EBNA-1 minus the repeat sequences (Figure 1) were synthesized by Sino Biologicals (Table S2). Cross-reactive peptides were synthesized by Alan Scientific (Table S2).

### Antibody Isotypes and Fc-Receptor Binding Activities Measurement

The level of antibodies and Fc-receptor binding activities were measured based on multiplexing Luminex microsphere-based method ^36,37^. Luminex beads were coupled to EBNA-1 peptides or cross-reactive peptides using a previously described method ^18^. Briefly, the carboxylate microspheres (Luminex) were conjugated covalently with the antigens (Key resource table) via NHS-ester linkages after activation from co-incubation with Sulfo-HNS and EDC (ThermoFisher). Sera were diluted at 1:200 for IgG1 and IgG3, 1:25 for IgG2, IgG4, IgM, and IgA, and 1:400 for Fc gamma receptors IIA and IIIA (FcγRIIA and FcγRIIIA). The samples were incubated with coupled microspheres in 384-well plates for 16 hrs. at 4°C, shaking at 750 rpm, to form immune-complexES. Then, the plates were washed with assay buffer (0.1% BSA and 0.02% Tween 20 in PBS) 3 times and incubated with PE-conjugated mouse anti-human antibodies for 1 hour at room temperature, at 750 rpm. Then, the plates were washed with the assay buffer 3 times, and all the samples were acquired by iQue Screener Plus (Intellicyt/Sartorius), or IntelliFlex Luminex Dual-Report (Luminex), and the median fluorescence intensities (MFI) were measured. All samples were performed in technical duplicates and averaged for final output.

### Antibody-dependent complement deposition

Antibody-dependent complement deposition (ADCD) was conducted using a 384-well plate format to evaluate the functional activity of antibodies in plasma samples ^38^. A Multiplex bead mixture was prepared by adding 15 μL of individual EBNA-1 peptide antigen-coupled beads to 18 mL of Luminex assay buffer. The remaining steps, including plasma incubation, complement activation, C3 detection, and analysis on the (iQue, Intellicyt), were performed according to a previously published method^38^

### Antibody-dependent cellular phagocytosis

Antibody-dependent cellular phagocytosis by monocytes (ADCP) was conducted to assess the phagocytic activity of THP-1 cells in response to antigen-coated beads opsonized with antibodies from plasma samples, following a previously published method ^39^. Briefly, biotinylated antigens were conjugated to Neutravidin FTIC beads, incubated with diluted plasma, and then exposed to THP-1 cells. After incubation and washing, cells were fixed and analyzed using the (iQue, Intellicyt). The phagoscore is calculated by taking the mean fluorescence intensity (MFI) of the beads that have been engulfed by the cells and multiplying it by the percentage of cells that have taken up the beads out of the total cell population. This product is then divided by 10,000 to yield the final phagoscore. The phagoscore provides a measure of both the intensity of bead uptake and the proportion of cells involved in phagocytosis, offering a comprehensive assessment of phagocytic activity in the sample.

### Peptide blocking assay to evaluate EBNA-1 and alpha crystalline beta cross-reactivity

Plasma samples from individuals with CRYAB antibody levels above the median (MFI = 1979) were selected for the peptide blocking assay (IM cohort: 6 months (n=11), 1-year (n=44), Seropositive (n=25), Seronegative (n=10). Plasma samples were diluted 1:50 and treated overnight with 1 µM each of the EBNA-1 peptides (aa 377-449, 365-420, 393-448). Influenza HA was used as a standardizing control. Following incubation, CRYAB-coupled Luminex beads were added to the plasma samples and incubated for 2 hours. The samples were then washed three times with Luminex assay buffer. After washing, total IgG anti-human secondary antibody tagged with PE was added and incubated at room temperature for 1 hour. This was followed by three additional washes with Luminex assay buffer (PBS 1X, 0.01% BSA, and 0.05 % Tween-20). Binding was quantified using the xMAP INTELLIFLEX® System. ^32^

### Statistical analysis

All statistical analyses were performed using R software (version 4.4.1, 2024-06-14, ucrt). All graphical visualizations were done using ggplot2 (version 3.5.1)^40^ package. To compare antibody responses across different EBV infection stages, we used Wilcoxon paired comparisons, with p-values adjusted for multiple comparisons using the False Discovery Rate (FDR) correction. For comparisons between DRB1*1501 positive and DRB1*1501 negative individuals, or other HLA alleles of interest, the two-tailed Mann-Whitney U test (Wilcoxon rank-sum test) was employed, with FDR correction applied to p-values. Fold changes between the acute and 1-year time points were calculated and these fold changes were compared between DRB1*1501 positive and DRB1*1501 negative groups using the Mann-Whitney U test. The lower limit of detection (LLOD) is defined as the mean of EBV seronegative (n =10) values plus three times the standard deviation. Values below the LLOD were set to the lower limit of detection before analysis. Mixed-effects models were also used to analyze the interaction between EBV infection stage and DRB1*1501 status, with infection stage and DRB1*1501 status as fixed effects and sample ID as a random effect. Statistical significance was defined as adjusted p values p < 0.05, with significance levels denoted by stars: *** p < 0.001, ** p < 0.01, * p < 0.05.

## Author contributions

**Conceptualization**: KL, RPM, RP, SC; **Methodology**: KL, RPM, KKG, MM, RP, SC, MC; **Investigation:** KL, RPM, RB2,3, RB1, KKG.; **Writing – Original Draft:** KKG.; **Writing –Review & Editing**: KL, RPM, RP, SC, MC, MM, KKG, QW, RB2,3, and RB1; **Data Curation:** MM, KKG, RB2,3; **Formal Analysis:** KKG, MM, KL, RPM; **Software:** RPM, KKG, MM, QW; **Visualization:** KKG, MM, RPM, WJ and QW; **Funding Acquisition:** KL and RPM; **Resources:** KL and RPM; **Supervision:** KL, RPM

---

**Figure S1.**
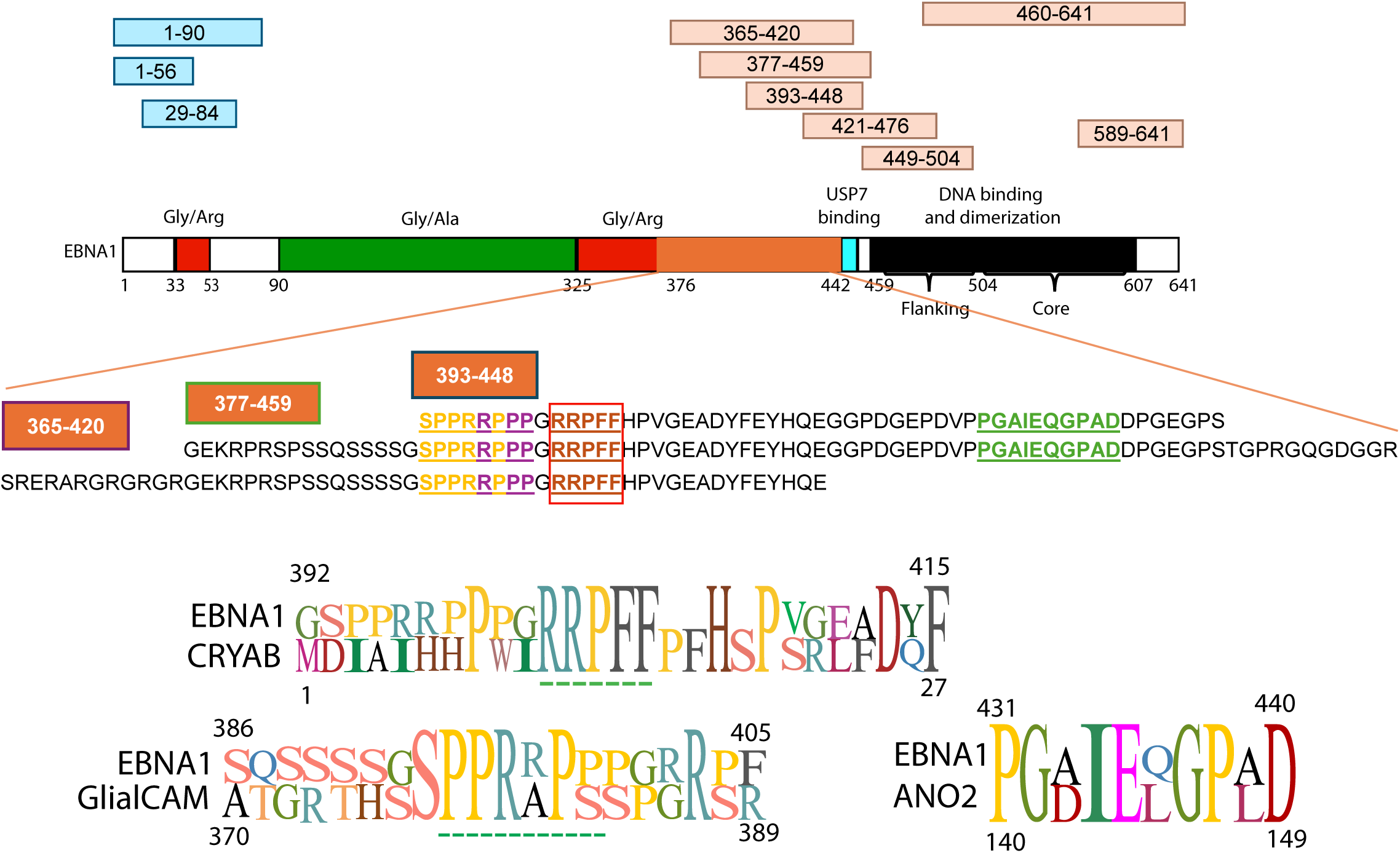
Schematic representation of EBNA-1 and homology with host proteins. The figure shows the EBNA-1 protein with regions highlighted for N-terminal and C-terminal peptides. Specific peptides (365-420, 377-459, 393-448, 421-476, 449-504, and 460-641) within the N- and C-terminal domains are highlighted in beige above the linear structure of EBNA-1. The lower section highlights regions where homology between EBNA-1 and host proteins is observed. Homologous regions between EBNA-1 peptides and host proteins GlialCAM (residues 370-389), CRYAB (residues 1-27), and ANO-2 (residues 140-149) are shown. Notably, sequences between residues 377-459 share homology with CNS proteins, including GlialCAM and CRYAB, with a specific overlap of the EBNA-1 C-terminal peptide region 393-448 and CRYAB.

**Figure S2.**
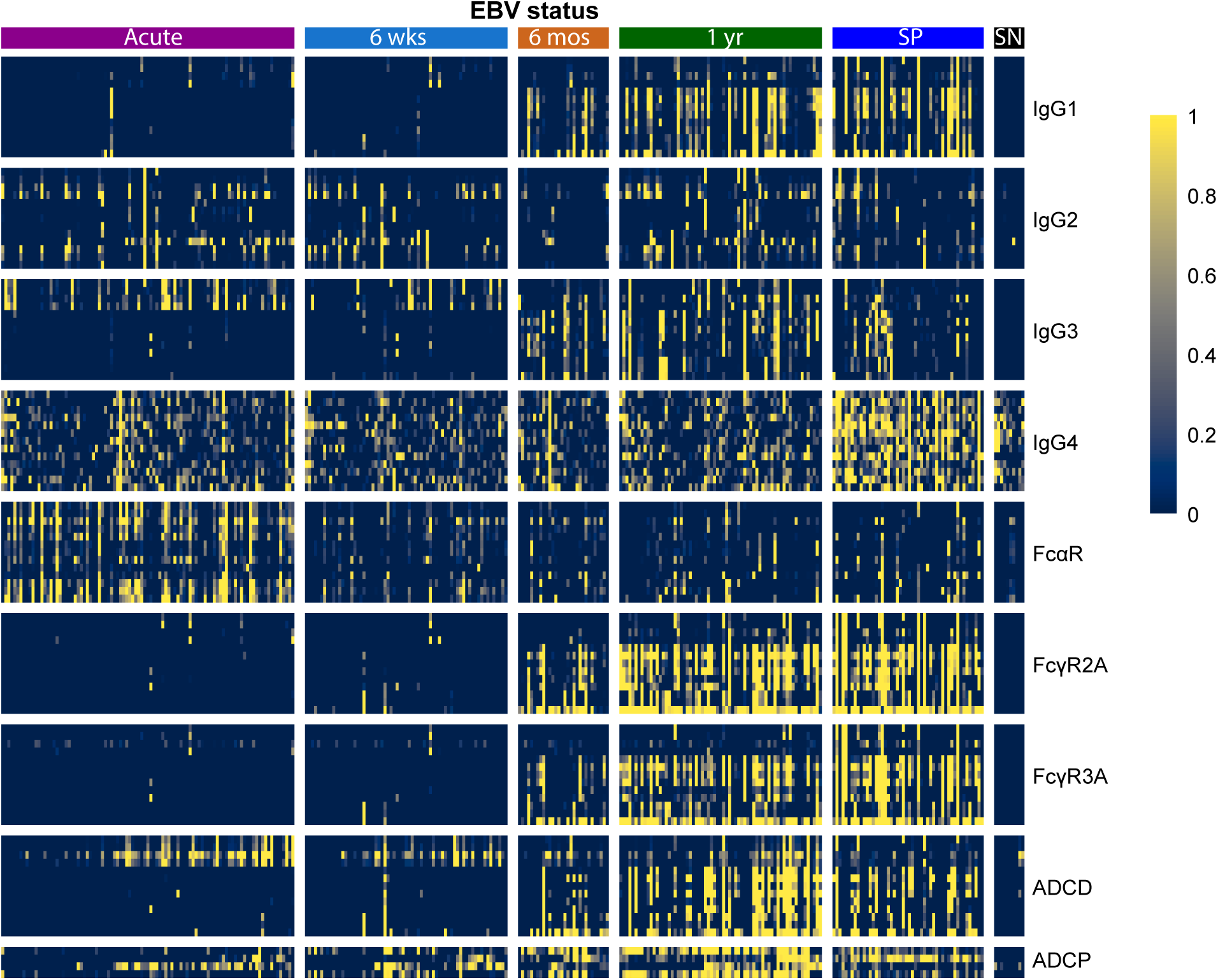
Heatmap of Normalized Immune Responses Across EBV-Specific Variables. Heatmap illustrating the normalized immune responses to various EBNA-1 peptides across different antibody isotypes and subclasses. This normalization highlights the relative magnitude of responses across groups while preserving the distinct profiles of each immune category. Data are stratified by EBV infection status, Acute presentation (n=97), 6 weeks (n=67), 6 months (n=30), and 1 year (n=67) post-IM diagnosis. EBV seropositive (SP, n=50) and EBV seronegative controls (SN, n=10). This visualization enables the comparison of response intensities across groups and categories, leading to the identification of IgG1 and IgG3 as focal points for subsequent analyses.

**Figure S3.**
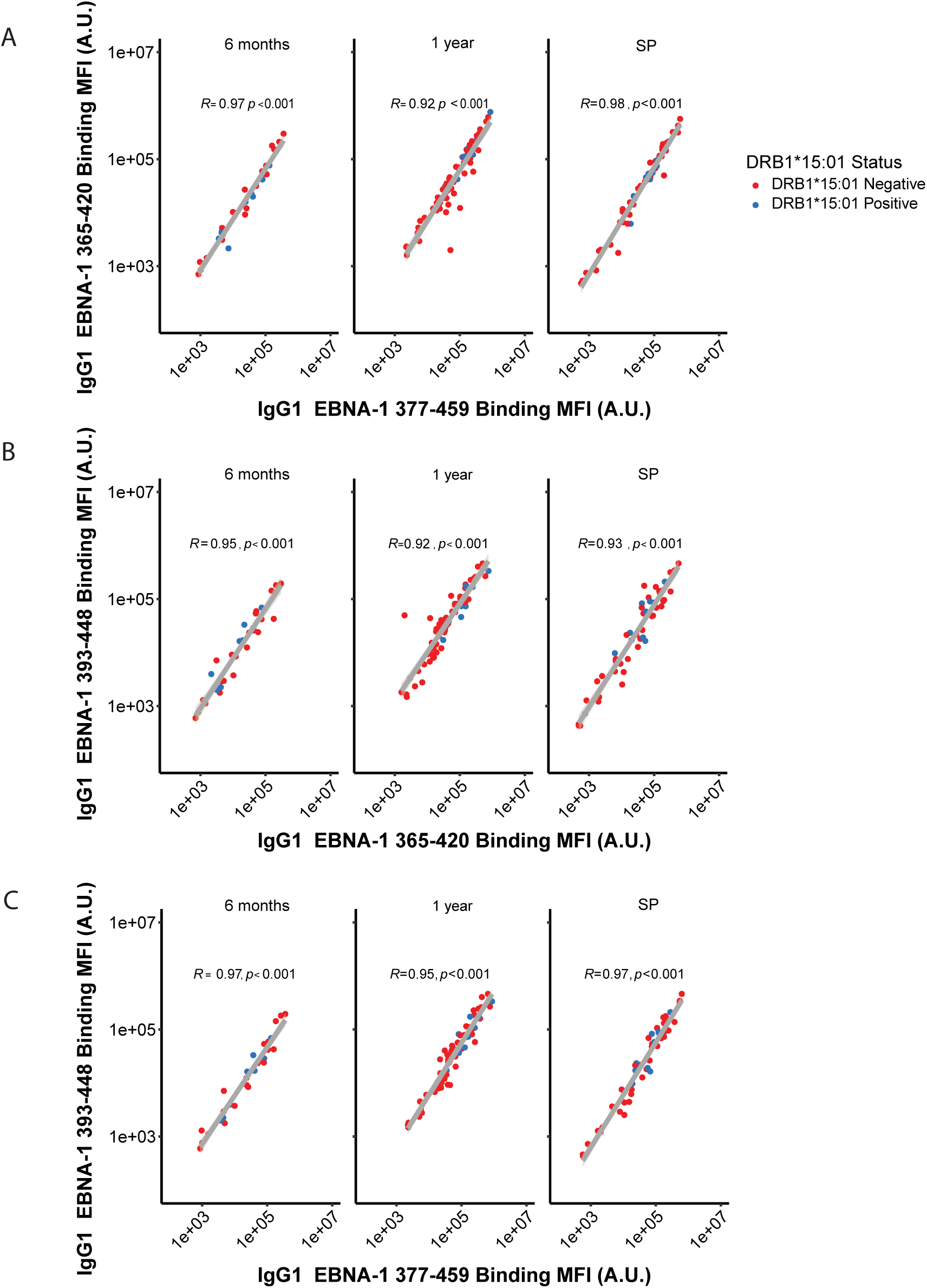
IgG1 binding correlation by DRB1*15:01 status. Spearman correlation analysis of IgG1 levels targeting C-terminal EBNA-1 peptides. Panels A-C represent pairwise correlations for IgG1 MFI values between peptides 365–420, 377–459, and 393–448, highlighting consistently high correlation levels at 6 months, 1 year, and in seropositive (SP) individuals. Data indicate robust cross-reactive responses among C-terminal EBNA-1 epitopes across infection statuses, with correlation coefficients (R > 0.9) and significance levels (p < 0.001) supporting consistent immune recognition patterns. Red and blue data points differentiate DRB1*15:01-negative and DRB1*15:01-positive individuals, respectively.

**Figure S4.**
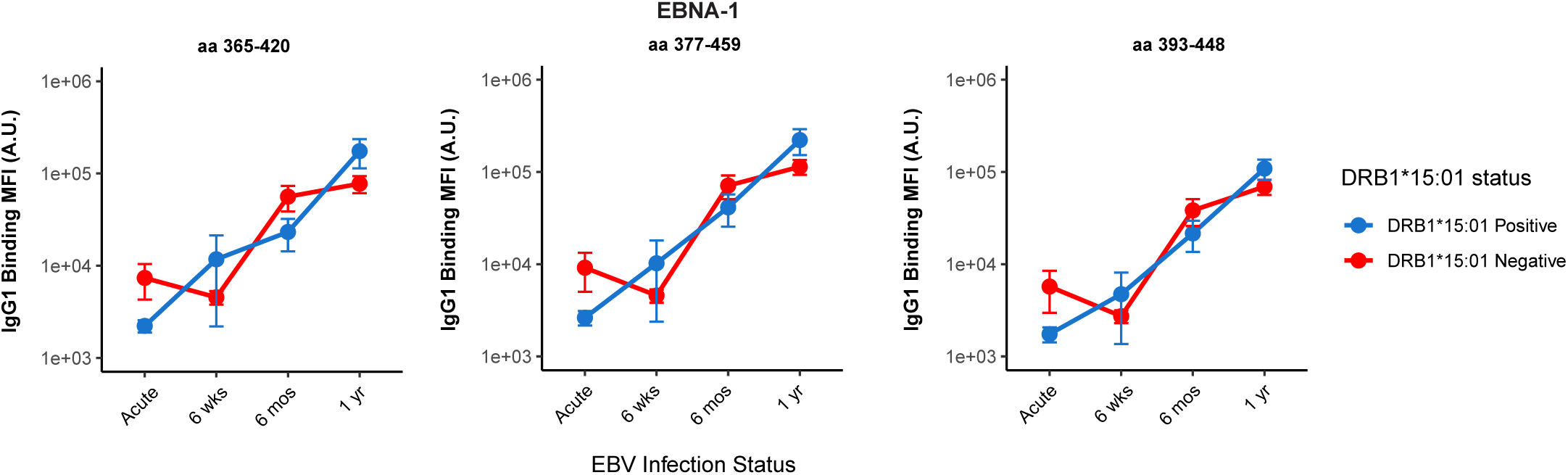
Longitudinal analysis of IgG1 binding to EBNA-1 peptides by HLA DRB1*15:01 Status. Line plots illustrate the trend in IgG1 binding to three EBNA-1 C-terminal peptides (377-459, 365-420, 393-448) across different infection stages (Acute IM, 6 weeks, 6 months, and 1 year). Data are stratified by DRB1*15:01 status, with red representing individuals without DRB1*15:01 and blue representing individuals with an HLA DRB1*15:01 allele. Error bars denote the standard error of the mean. A mixed model test indicates significant effects of infection stage and DRB1*15:01 status on antibody binding trends. Individuals with at least one DRB1*1501 allele exhibited consistently higher binding responses over time, with significant DRB11501 influence noted at 1 year for aa377-459 (p=0.003), aa365-420 (p=0.001), and aa393-448 (p=0.05).

**Figure S5.**
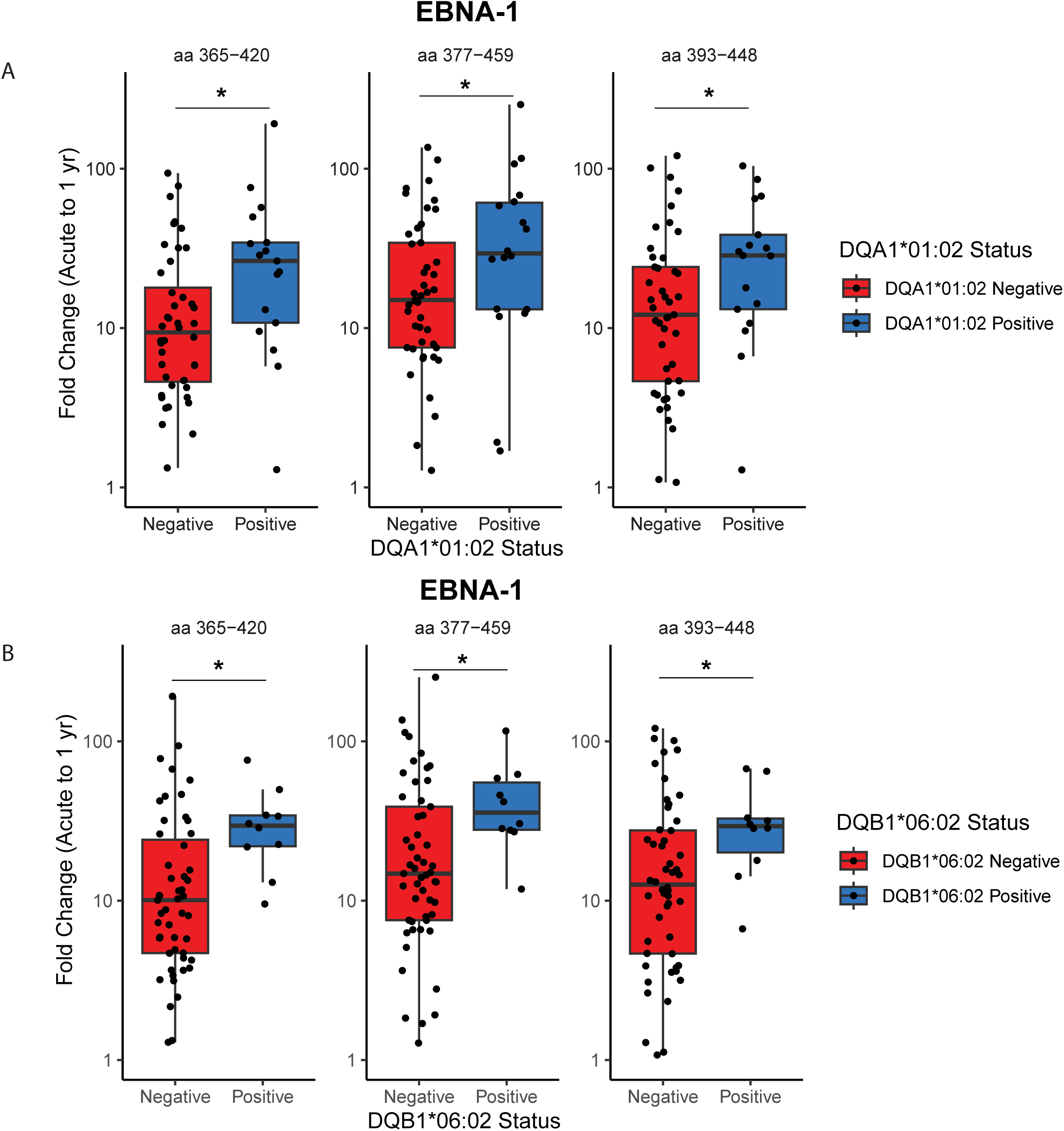
Fold-change comparisons of EBNA-1 IgG1 responses by DQA1*01:02 and DQB1*06:02 status. This figure illustrates the fold change in IgG1 binding responses to EBNA-1 C-terminal peptides (aa 365-420, aa 377-459, and aa 393-448) from acute IM to 1-year post-infection, stratified by HLA alleles. Panel A shows individuals with at least one DQA101:02 allele. Red indicates individuals without DQA101:02-negative, while blue denotes individuals with the DQA101:02 allele. Significantly higher fold change is observed in individuals with DQA101:02 compared to those without this HLA allele across all peptides (*p < 0.05). Panel B shows higher fold changes in EBNA-1 IgG1 binding antibody levels in individuals with DQB106:02 compared to levels in those without this allele (*p < 0.05). Statistical significance is denoted by asterisks: *p < 0.05.

**Figure S6.**
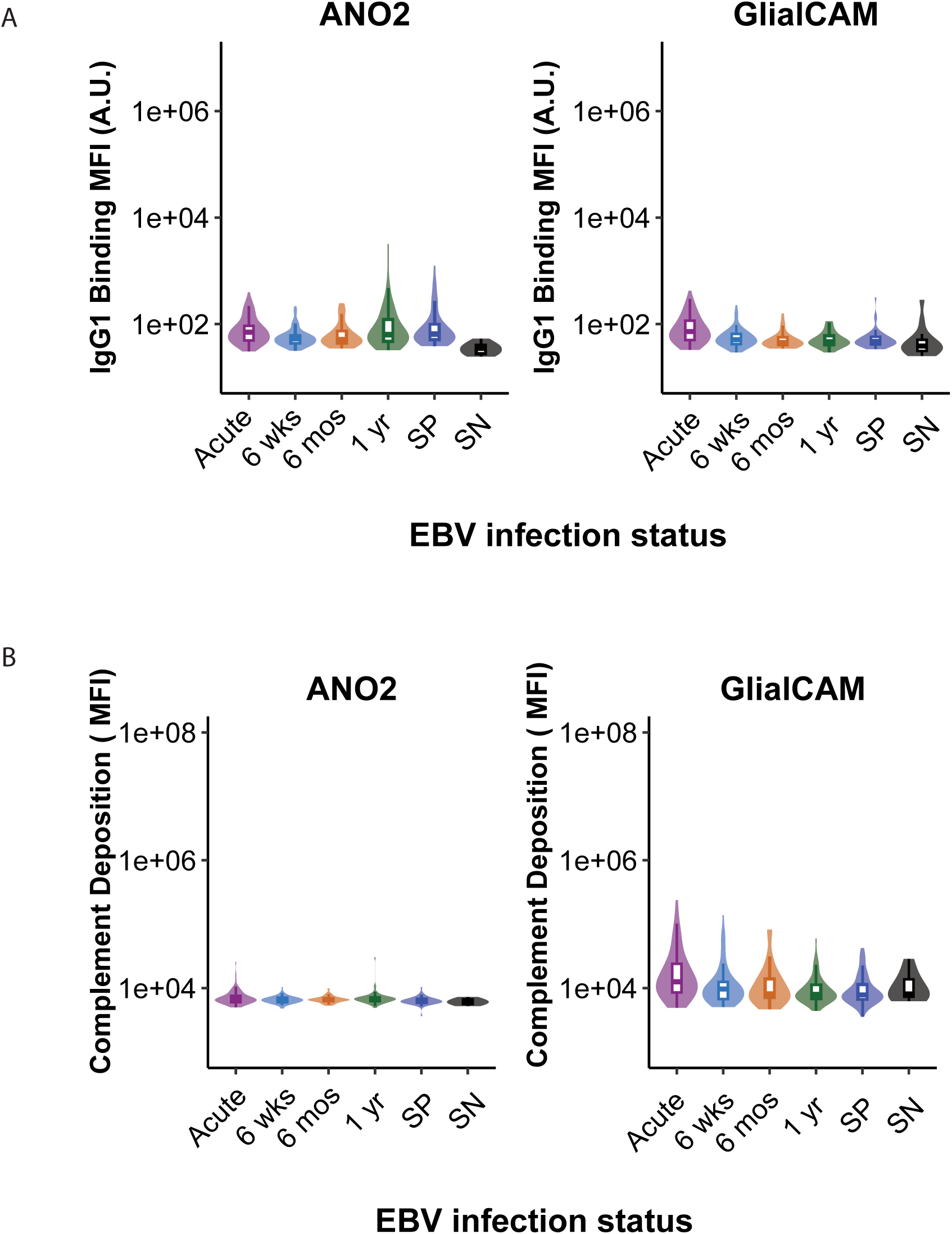
IgG1 binding and ADCD for Anoctamin and GlialCAM. This figure demonstrates a lack of IgG1 binding and complement deposition against self-antigens anoctamin (ANO2) and GlialCAM, across different stages of EBV infection. Panel A: IgG1 binding MFI (arbitrary units) to ANO2 and GlialCAM is shown across Acute, 6 weeks, 6 months, 1 year, SP (Sero_Pos), and SN (Sero_Neg) groups. Panel B: Complement deposition (MFI) levels for ANO2 and GlialCAM similarly show negligible responses, with consistent low-level deposition across all stages, underscoring the specificity of observed immune responses to EBNA1 and CRYAB but not to these non-cross-reactive proteins.

